# A practical sampling strategy for biodiversity genomics of reptiles: The *Cryptoblepharus pulcher* assembly offers insights into the demise of a threatened relative

**DOI:** 10.64898/2026.08.27.747638

**Authors:** Tristram O. Dodge, Mario Ernst, Paul Oliver, Mozes P.K. Blom

**Affiliations:** Dept. of Biology, Stanford University, Stanford, USA; Dept. for Evolutionary Diversity Dynamics, Museum für Naturkunde Berlin, Leibniz Institute for Evolutionary and Biodiversity Science, Berlin, Germany; Institute for Biology, Humboldt-Universität zu Berlin, Berlin, Germany; Centre for Planetary Health & Food Security, Griffith University, Nathan, Australia; Queensland Museum, South Brisbane, Australia

**Keywords:** Biodiversity genomics, long-read sequencing, reference genome, skink, Cryptoblepharus

## Abstract

Due to the sparse and uneven availability of genomic resources, it remains challenging to appraise genomic attributes for species of conservation concern. While long-read sequencing enables assessment of genetic diversity at unprecedented scale, accessing high-quality tissues remains a challenge for non-model species. Here, we explore an alternative sampling strategy for tissues where “gold-standard” cryopreservation is infeasible. Focusing on the Australian scincid lizard *Cryptoblepharus pulcher*, we compare DNA obtained from various ethanol-preserved tissue types and DNA extraction kits, and ask whether high-molecular weight DNA can still be retrieved. PacBio HiFi sequencing of the most promising sample yielded a highly contiguous reference-level assembly, validating this approach in vertebrates, specifically lizards. After scaffolding the assembly to chromosome-level, we then used a comparative approach to shed light on the evolution and demise of *C. egeriae*, a closely related, now Extinct-in-the-Wild species. Surprisingly, despite being a wide-spread continental analogue with a similar ecology, *C. pulcher* has lower genetic diversity and long-term historical population size than *C. egeriae,* an island endemic. However, *C. pulcher* also shows fewer runs-of-homozygosity, supporting prior reports that *C. egeriae* experienced recent inbreeding. Together, these findings demonstrate that a practical and cost-effective preservation strategy can still yield high-quality genomic resources in vertebrates, as well as valuable insights that are relevant in an age of biodiversity decline.

## Background

Understanding the genetic context of biological diversification and decline is of fundamental importance for both evolutionary and conservation research. High-quality genomic resources are a key requisite to facilitate such studies [1]. The increasing availability of reference genomes has already provided unique insights into genome evolution, diversity and architecture, and how this translates into the evolutionary mechanisms shaping biodiversity [2]. While the assembly of high-quality reference genomes was historically a major challenge, advances in long-read sequencing (LRS) have substantially lowered the technical and financial needs to produce a reference genome for many organismal groups. Yet, the availability of high-quality genomic resources is still highly uneven across eukaryotes [3] and the number of reference genomes for major taxa does not scale with taxonomic or evolutionary diversity. Even among amniotes, arguably the most well studied eukaryotic clade from a genomic perspective, genomic research has primarily focused on avian [4] and mammalian groups [5].

In contrast, Squamates (lizards and snakes) are the most species rich and evolutionarily diverse group of amniotes, but taxonomic sampling is still so sparse that in many instances there is only one or, at most, a few species per family with genome-scale data [6]. While the number of squamate genomes is on the rise, the unavailability of genomic data for closely related taxa still means that we often lack sufficient resources to undertake finer scale comparative analyses of genome structure and evolution. This hampers our ability to understand the genetic underpinnings of major evolutionary transitions and to determine the extent to which genomic attributes of reference species are idiosyncratic or shared across lineages [6]. For example, *Anolis carolinensis* was for many years the only squamate genome available [7] and its genomic characteristics such as genetic diversity, repeat landscape and the distribution of GC-content were considered as archetypical for Iguanids and other lizards. However, sequencing additional species [8,9] has shown that these estimates vary among *Anolis* species and that the reference genome of *A. carolinensis* may be rather unusual due to multiple genetic bottlenecks during a recent expansion from Cuba and Florida [10]. Such examples emphasize the importance and the need for producing a more balanced distribution of genomic resources across eukaryotes. To achieve this objective there is a growing demand for high-quality tissue samples that are suited for LRS.

The success of LRS strongly scales with DNA quality (continuous integrity of the DNA molecule) but preserving DNA quality is not a trivial exercise when working with non-model organisms. The ‘gold-standard’ for DNA preservation (see for overview [11,12]) includes cryopreservation for sampling and transfer, but this is often extremely costly and logistically challenging; particularly in remote locations. Yet, most biodiversity hotspots tend to be located in remote regions or heterogeneous landscapes that are difficult to access [13]. Alternative protocols have been proposed and thoroughly tested for well-studied taxonomic groups (e.g. birds: Queen’s buffer [14], or mosquitoes: EtOH [15]), but for most taxa the efficacy of cost-effective and practical substitutes remains largely unexplored. As a consequence, access to high-quality tissue samples is a major concern for many projects aiming to generate reference genomes or to assay all aspects of genetic diversity at a population level scale (e.g. pangenomes; [16]). Moreover, with biodiversity declining at an unprecedented rate, the paucity in high-quality samples from natural populations is likely to expand and will have major consequences for our ability to assay the genetic diversity underlying contemporary biodiversity. We have therefore recently called for a concerted effort to develop and test explicit methods that preserve DNA integrity across taxonomic groups and work towards the expansion of high-quality biodiversity tissue archives [11].

Where others (e.g. [12]) have contributed to this objective by presenting findings based on standardized laboratory tests, here we complement such approaches by mimicking a typical fieldwork-to-lab workflow in an understudied reptile genus where some species have recently been reported as Extinct-in-the-Wild.

Lizards of the genus *Cryptoblepharus* are small (∼3-6 cm, snout-vent length [SVL]) diurnal skinks that originated in South-East Asia, rapidly radiated across Australia, and naturally dispersed across both the Pacific and Indian Ocean [17]. Within Australia, they diversified in two major clades and represent a continental adaptive radiation [18,19]. While most species are relatively common, the Christmas Island endemic (*C. egeriae*) was declared as Extinct-in-the-Wild in 2017 [20]. This species now primarily persists in captive breeding colonies [21], although a small number of captive bred individuals have now been released and are actively monitored on two islands of the Cocos (Keeling) atoll [22]. A high-quality *C. egeriae* reference genome was recently assembled with PacBio HiFi reads and subsequently used to assess evolutionary history and recent demography [23]. Interestingly, despite being an island endemic, high genome-wide heterozygosity suggested that the *C. egeriae* population used to be large but long runs of homozygosity (ROH) also hinted that recent inbreeding may have taken place before or after the onset of captive breeding. However, given a lack of genomic resources within Scincidae, the magnitude of these observations could only be compared to very distantly related organismal groups that differ radically in terms of ecology, life-history and, likely, the rate of genome evolution; all factors that shape genetic diversity.

Thus, a high-quality genome assembly for a congeneric species is needed to provide a comparative window on genomic and demographic change in *Cryptoblepharus* species. *C. pulcher* is a medium sized (∼4 cm SVL) *Cryptoblepharus* that can inhabit a variety of tropical to temperate woodland and open forest habitat [24,25]. It is an arboreal ecomorph that is frequently found in forest habitat along coastal eastern and southern Australia [24]. Among continental *Cryptoblepharus, C. pulcher* putatively matches the habitat requirements and ecological niche of *C. egeriae* [18,24] most closely.

Here, as a proof of concept, we employ a practical fieldwork-to-lab sampling, storage, and transfer strategy for lizard samples. We compare sample quality across different tissue types and extraction protocols. We subsequently generate Pacific Biosciences (PacBio) High-Fidelity (‘HiFi’) reads for the most promising extract and assemble a high-quality reference genome. We used data from our focal taxon—the lizard *C. pulcher*—to provide a comparative genomic view on both the demographic history and genetic composition of a congeneric species which recently went extinct in the wild, *C. egeriae*.

## Methods

### Sampling strategy - LRS

From May to June 2021, we collected eight *C. pulcher* individuals by hand in woodland habitat (-27,5383; 153,044) near Brisbane, Australia. Individuals were kept alive and brought into the lab where they were euthanized shortly after. Rather than using liquid nitrogen for snap-freezing samples and cryopreservation (the ‘gold standard’ for DNA preservation [12]), individuals were dissected at ambient temperature and tissues were directly transferred to cooled down tubes (4 °C) filled with >95% EtOH (Sigma-Aldrich; #E7148). For each individual, we aimed to collect four different tissue types: liver, heart, gonads and muscle. However, due to the small size of the species, it sometimes can be challenging to sample all organs, and it is often unavoidable that other tissue types may still be attached when targeting specific tissues. For example, it is likely that surrounding soft tissue may have been excised when sampling the gonads. Each tissue was sampled as fast as possible following euthanasia and small incisions were made with a scalpel to promote rapid penetration of EtOH within tissues. Following tissue collection, tissues were stored up to five days in a fridge (4 °C) rather than a freezer to minimize freeze/thaw cycles prior to shipping.

To preserve DNA molecule integrity, courier shipping tissue samples on dry ice is ideal but it comes with substantial costs and bears a risk if shipping is delayed and dry ice top-up is not feasible. Instead, we shipped samples for LRS with regular post (Australia Post to DHL), without cooling, and the tissues were shipped in screw-cap tubes filled with >95% EtOH. For this study, we selected >95% EtOH as primary storage solution because i) it has been shown to work well at room temperature for other taxa [15,26], ii) it is easy and affordable to obtain, and iii) it can be shipped in small quantities under IATA Special Provision A180. The packaging of samples was done in compliance with the rules and regulations as outlined in IATA Special Provision A180. Total transfer time between the lab in Brisbane (Australia) and the lab in Berlin (Germany) was 12 days. Upon arrival, all samples were immediately stored at 4 °C, and DNA extractions were completed within a week.

### Extraction comparison

The efficacy of LRS depends on both DNA quality (the length of the DNA fragment) and the purity of the DNA extract. Furthermore, for small organisms such as *Cryptoblepharus*, another parameter of importance is the amount of DNA that can be retrieved from a single extraction. Ideally a single extraction yields enough DNA for library preparation and avoids the need for in-vitro DNA amplification. Yet, while some tissues may yield a lot of DNA, the quality or purity of extracts can differ between tissues. For example, it has been suggested that liver tissues yield more degraded DNA than other tissues due to enzymatic activity and rapid post-mortem autolysis [27]. We explored the suitability of four commercial DNA extraction kits specifically tailored for obtaining High-Molecular Weight (HMW) DNA (see for overview of kits and catalog numbers: Supp. Table 1). For each extraction method we followed the manufacturer’s protocol, conducted a DNA extraction with similar sized tissue samples, for at least three different tissue types, and measured the quantity, quality and purity of extracts. DNA quantity was estimated using a Qubit fluorometer (Thermo Scientific). DNA quality was estimated by measuring the amount of HMW DNA (here characterized as % of fragments >40 kilobases (kb) in length) using a TapeStation (Agilent). The purity of extracts was assessed by using the A260/280 and the A260/230 absorbance ratios, estimated with a NanoDrop spectrophotometer (NanoDrop Technologies). Following an initial screen using all extraction kits and tissue types, we then repeated extractions for kits and tissues that seemed most promising (gonad/liver and the Circulomics kit). All DNA extracts were eventually stored at -80 °C. The most promising extraction (Circulomics kit), based on quality, quantity and purity measurements, was from a gonad tissue from a single individual. This DNA extract was shipped (<12 hours) on dry ice from Berlin to the Science for Life Laboratory (SciLifeLab) in Uppsala (Sweden).

### PacBio HiFi library preparation and sequencing

Library preparation, quality control and sequencing were all done at SciLifeLab. Briefly, HMW DNA was sheared to 15-20 kb fragments using a Megaruptor 3 (Diagenode), and size selection was done with a SageELF (Sage Science) system. The resulting DNA extracts were used as input to the SMRTbell Express Template Prep kit 2.0 to prepare libraries suitable for sequencing on the PacBio Sequel II platform. AMPure Beads (Beckman Coulter) were used for clean-ups. Sequencing was done with two single-molecule real-time (SMRT) 8M cells.

### Genome assembly

HiFi reads (read quality >99) were selected from the circular consensus sequencing (CCS) UBAM with bamtools v2.5.1 [28] and reads that contained adapter sequences were filtered using HiFiAdapterFilt v3.0.1 [29], using as parameters 30 length and 97% match. Read quality of the filtered dataset was assessed using NanoPlot v1.42 [30].

To assemble the *C. pulcher* mitochondrial genome, we first ran mitoHiFi v2.2 [31] on the filtered HiFi reads, specifying the *C. egeriae* mitochondrion as the reference sequence (NCBI Accession: CM057396). The preliminary mitochondrial genome was a 31 kb sequence, roughly twice the expected length of the *C. pulcher* mitochondrion. To investigate and correct the sequence, we performed an alignment to other *Cryptoblepharus* mitochondrial genomes [32] and trimmed the sequence that aligned poorly, resulting in a 15,613 bp sequence (deposited on NCBI: CM145273).

The nuclear genome was assembled with hifiasm v0.16.1-r375 [33] using default parameters. An initial assessment of depth of coverage of HiFi reads mapped to the draft genome revealed several > 1 Megabase (Mb) regions where read depth was approximately half of the genome-wide average, indicating the possible presence of several erroneous duplications. We therefore ran purge_dups v0.0.3 [34] with default parameters on the primary hifiasm contigs.

### Omni-C library preparation and sequencing

As a proof-of-concept, the pragmatic sampling approach already yielded a high-quality *C. pulcher* assembly. However, we then opted to improve contiguity by using chromosome conformation capture and scaffold contigs into chromosome models. To this end, a second round of samples (seven *C. pulcher* individuals) was collected from the same locality in March 2022. In contrast to the samples collected for PacBio HiFi sequencing, the samples collected for Omni-C were snap frozen and stored in a -80 °C freezer prior to shipping with dry ice. They were shipped and transferred within 48 hours on dry ice as carry-on luggage during a commercial flight. The Dovetail Omni-C (Dovetail Genomics) kit and Illumina short-read sequencing were used to generate uniform chromatin conformation capture data at SciLifeLab Stockholm (Sweden). The Omni-C library was created, following the manufacturer’s protocol, using flash frozen liver tissue from a single individual. The Omni-C library was sequenced with a ¼ lane on the Illumina NovaSeq 6000 platform, generating 1.19 billion paired-end reads.

### Scaffolding the C. pulcher genome

To achieve a chromosome-level assembly, the filtered contigs were scaffolded into linkage groups using the Omni-C data collected from the additional *C. pulcher* individual. The Omni-C reads were mapped to the filtered contigs using the Arima Genomics mapping pipeline (A160156 v02) and we used the resulting BAM file (sorted by name) as input to the scaffolding tool YaHS v1.2a.2 [35]. We visually assessed the contact matrix in Juicer v1.1 [36] and made 1 manual join of a small contig to scaffold 2 based on contact density. We sorted and named scaffolds by length. Assembly contiguity was assessed using seqkit stat v2.10.0 [37] and assembly completeness evaluated with BUSCO v5.3.2 [38] using the vertebrata_odb10 dataset.

### Scaffolding the C. egeriae genome

The previously assembled high-quality *C. egeriae* genome, based on cryopreserved tissue samples, was generated using PacBio HiFi data, but no Omni-C data was available at the time for further scaffolding [23]. Nonetheless, the HiFi data alone already yielded 10 contigs that appeared to be complete telomere-to-telomere (T2T) chromosome models, while other chromosomes were likely broken into several contigs. To further improve and evaluate the *C. egeriae* assembly, we scaffolded the *C. egeriae* contigs to the empirically informed *C. pulcher* chromosome models using RagTag v2.1.0 [39]. The mapping parameter was modified to -x asm20, to account for the divergence between species. We investigated the AGP file and corrected two misjoins of short contigs to the telomeric ends of complete chromosomes to also produce the final scaffolded *C. egeriae* assembly.

### Repeat annotation and genome synteny

To annotate repeats in this non-model species, we built a repeat library with RepeatModeler v2.0.3 [40]. After six rounds of modelling, the resulting consensus library was used as input for RepeatMasker v4.1.2 (http://www.repeatmasker.org) to identify and mask repetitive sequences in the genome. Telomeric repeats were identified using the telo command in seqtk v1.4-r130 (https://github.com/lh3/seqtk).

To evaluate synteny between *C. pulcher* and *C. egeriae*, we performed a whole-genome alignment using minimap2 v2.24 [41] and the option -x asm20. The 15 longest sequences, representing the putative chromosomes, were aligned and the resulting PAF file was filtered to exclude alignments less than 1 Mb. We visualized these alignments using the package Circlize v0.4.16 [42] in R v4.4.1. To investigate synteny between the putative sex chromosomes, alignments were generated for each chromosome pair using minimap2 with the same parameters as above and the results plotted using custom R scripts.

### Identifying sex-linked sequences

Previous research on sex chromosomes in skinks [43] including *Cryptoblepharus* [23] indicated that the XY sex chromosome pair of *C. pulcher* would most likely be homomorphic. To identify the sex chromosome, we first assessed read-depth across the 16 chromosomes. Minimap2 was used with the options -x map-hifi and --secondary=no to map HiFi reads to the genome, and alignments were kept with mapping quality > 20. We then calculated mean read-depth in non-overlapping 1 Mb windows by using bedtools v2.30.0 genomecov [44]. Coverage was relatively consistent across the genome, but we identified two scaffolds (7 and 16) that displayed regions with mean read depth of half the genome-wide average (Figure S2). To investigate if these sequences might represent divergent alleles linked to the sex chromosomes (i.e. a possible XY pair), we first searched for signals of shared homology between scaffold 16 and other chromosomes, by surveying changes in coverage when mapping reads to a reference with and without scaffold 16. Read-depth across the genome remained largely consistent between analyses, with two exceptions: With scaffold 16 excluded, there is a higher coverage on scaffold 7 (within the previously identified region of ∼half depth), as well as within a 1 Mb window on scaffold 5 (Figure S2). The observed pattern of read sharing between scaffold 16 and scaffold 7, but rarely between scaffold 16 and other chromosomes, is consistent with distant homology between these sequences and supports a model where they may represent an old sex chromosome pair.

As orthogonal evidence, we also investigated Omni-C contact density and found that scaffold 16 showed strong affinity with the sequences flanking the half coverage region (i.e. the putative pseudoautosomal regions of scaffold 7) but not with the X and Y specific regions. This suggests that scaffold 16 represents a partially assembled alternate haplotype at this locus, without containing the pseudoautosomal sequence (Figure S3) which is assembled as part of scaffold 7.

To confirm that the shorter scaffold represented a partial Y chromosome and the longer scaffold represented the X chromosome, blastn v2.16.0 [45] was used to search for known X-linked genes shared across Scincidae [43]. We found that these genes only blasted to scaffold 7 and not to scaffold 16, consistent with patterns in *C. egeriae* (Dodge et al., 2025). Finally, we performed whole chromosome alignments of these scaffolds with the *C. egeriae* genome. We found that both the X alignments and Y alignments shared sequence identity. Thus, several observations suggest that the 7th longest scaffold in our *C. pulcher* assembly represents a near-complete X chromosome, while the 16th longest likely represents the sex-linked region of the *C. pulcher* Y chromosome. Based on these observations, we renamed the latter sequence chrY and treated it as a partial Y chromosome in subsequent analysis.

### Mapping and variant calling for population genetic analyses

Our population genomic analyses relied on mapping reads to their respective reference genomes. However, correctly representing both sex chromosomes in a genome is not trivial, in part because this region of the genome is diploid in an otherwise haploid reference. As such, we mapped reads to the complete reference genomes for each species and repeated the procedure with chrY omitted. Minimap2 was used to map the HiFi reads with the options -x map-hifi and --secondary=no to only include primary alignments. We filtered the resulting BAM file to only include reads with mapping quality > 20. To map Omni-C reads, we again followed the practices laid out by the Arima Genomics mapping pipeline. Briefly, the Omni-C library was paired-end; however, because insert sizes vary dramatically for Omni-C data and can even be inter-chromosomal, we mapped each ∼150 bp read as single-end. We then used custom scripts to combine the reads into a single BAM file, mark and filter duplicates, and only keep reads with mapping quality > 20.

Variants were called using GATK v4.6.0.0 [46] for both the HiFi and Omni-C data. We used HaplotypeCaller with -pcr-indel-model AGGRESSIVE to generate an initial GVCF and then ran GenotypeGVCFs with the --all-sites parameter to generate an all-sites VCF file. We followed GATK best practices for variant filtration, with slight modifications that have been shown to have good performance in *Cryptoblepharus* [23], namely a more stringent quality by depth (QD) filter of 10 for single nucleotide polymorphisms (SNPs) and insertions/deletions (INDELs), and requiring a reference genotype quality (RGQ) value of 20 for INVARIANT sites. Because read depth varied across libraries, we also employed depth filters based on mean read depth for each data type, requiring a region to be covered by at least 10 reads and not exceed 1.5x mean depth. For the *C. pulcher* HiFi and Omni-C data, our upper depth threshold was 40x and 110x, respectively, and was 64x for the *C. egeriae* HiFi data. For computational efficiency, we parallelized each step by chromosome and later combined the filtered VCFs.

### Inferring historical population sizes and inbreeding estimation

We assembled the *C. pulcher* genome to expand our perspective on the demographic estimates and trajectory inferred for *C. egeriae* [23]. We therefore primarily aimed to calculate genetic diversity (heterozygosity), assess the extent of inbreeding using runs of homozygosity (ROH) and the change in effective population size over longer time scales (PSMC). However, we leveraged our sequencing strategy to calculate these attributes using two distinct data types generated from two different *C. pulcher* individuals: i) The long-read HiFi data and ii) the short-read Omni-C Illumina data. All downstream analyses were conducted in the same way, irrespective of data type.

Genome-wide estimates of heterozygosity were calculated using two different window sizes (10 kb and 1 Mb) with pixy v1.2.10 [47]. Heterozygosity was then visualized across the entire genome in 1 Mb bins. We quantified the abundance of long runs of homozygosity (ROH) as a proxy for the extent of inbreeding [48]. Long ROH are commonly defined as >1 Mb and we detected these using a custom R script find_ROH [23]. This implements an observational approach that has been shown to sometimes outperform other methods (e.g., plink --homozyg) in HiFi data from *Cryptoblepharus* [23]. Briefly, this script calls ROH by identifying consecutive 10 kb windows with < 2 heterozygous SNPs per window or if the two adjacent windows have fewer than 2 SNPs. As input, we used the pixy 10 kb file from the genome with the Y included (to avoid spurious mapping of Y-linked contigs to the autosomes breaking up ROH) but excluded windows with fewer than 5000 called sites. We note that the mean number of SNPs in these bins are 50.9 and 53.4 for the *C. pulcher* HiFi and Omni-C data, respectively, and 61.1 for the *C. egeriae* HiFi data. An analysis with plink v1.90b6.5 – homozyg [49] yielded similar results but, since several long ROHs appeared spuriously broken, we focus on the findings of our custom method. Finally, we filtered out ROH calls in the putative sex-linked region on chromosome 7, due to low confidence in hemizygous regions.

To infer the demographic trajectory over longer evolutionary timescales, we ran PSMC v0.6.5-r67 [50] which implements the Pairwise Sequential Markovian Coalescent method. Sex-chromosomes were excluded for this analysis, and we only retained autosomes in the VCF file. We initially used the default PSMC parameters of -N 25 -t 12 -r 5 -p “4 + 25*2 + 4 + 6”, and found low bootstrap support in recent time segments. We tested if this might be due to model misspecification, as has been recently reported [51], and modified the parameter -p “2 + 2 + 25*2 + 4 + 6” and -p “1 +1 + 1 + 1 + 25*2 + 4 + 6”. However, bootstrap support remained low and we therefore excluded the first 6 atomic time segments for each individual, due to low confidence at these time scales.

## Results

### DNA quality, quantity and purity: Tissues and Kits

To identify the most promising HMW DNA extraction strategy, we used at least three different tissue types (liver, gonad, muscle) across four extraction kits and used heart tissue when available (2x). *Cryptoblepharus* skinks are relatively small and slender lizards, and it was challenging to dissect the heart muscle for two other individuals. Due to the small size, the two extractions with heart tissue likely also yielded a very low quantity of DNA and may therefore not be an optimal tissue type for small lizards. Similarly, it was hard to obtain large amounts of muscle tissue and these samples also consistently yielded relatively low quantities of DNA; regardless of extraction kit (Figure 1).

**Figure 1.**
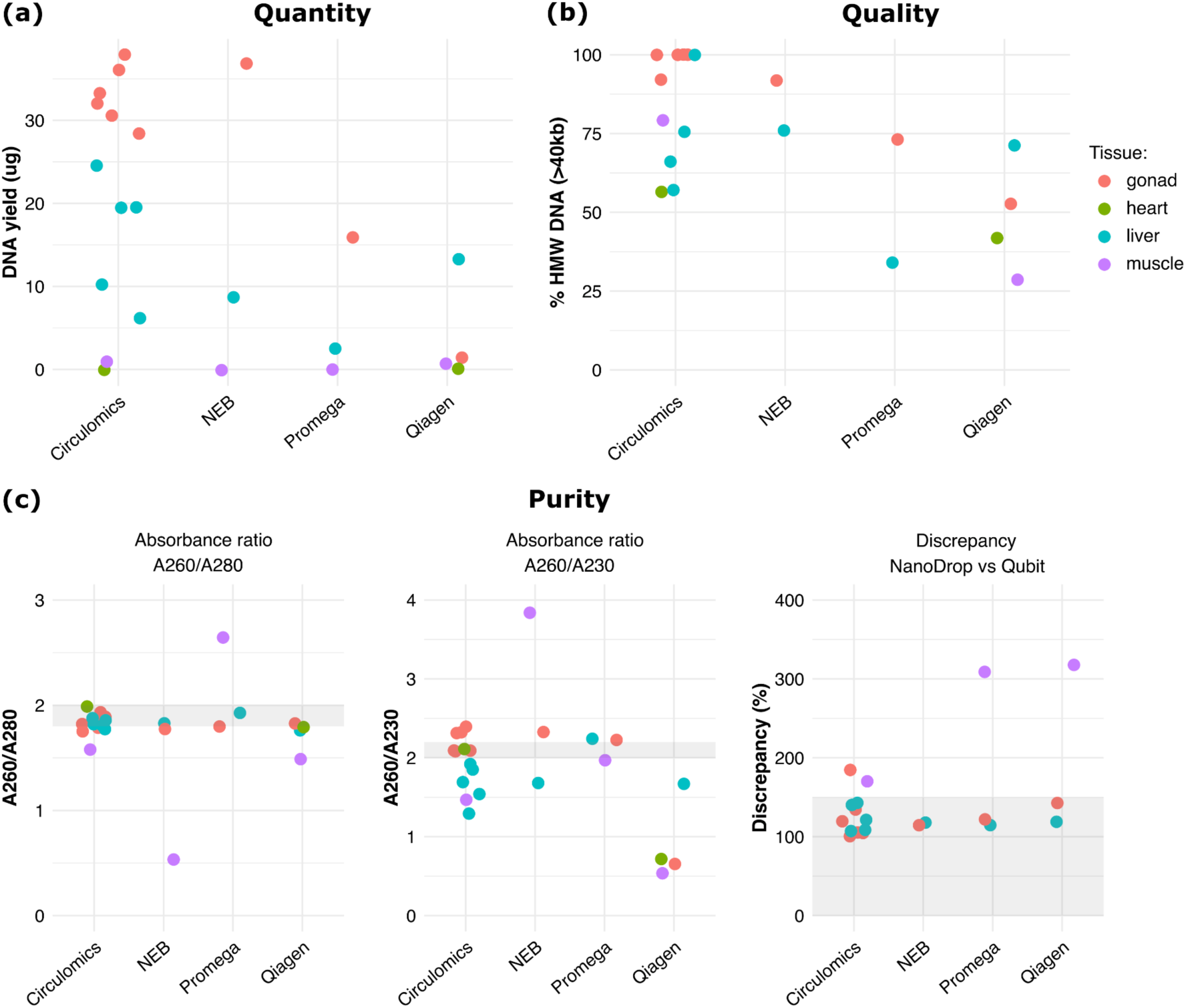
Comparative analyses of quantity, quality and purity of DNA extracts. Yield of DNA quantity (a), quality (b) and purity (c) across four different tissue types and DNA extraction kits. All tissues were in transit for 12 days at ambient temperature in >95% EtOH. DNA extracted with the Circulomics kit from gonad tissues consistently yielded more and better-quality DNA, at purity levels suited for PacBio HiFi sequencing, than other tissue or kit types. Gonad and liver extractions with the Circulomics kit were repeated to evaluate consistency and gonad tissues were less variable in terms of DNA yield and quantity.

Even though all tissues were sampled and transferred in the same way, gonad and liver tissues consistently yielded more DNA and a higher ratio of HMW DNA than the other tissue types (Figure 1). For two extraction kits (NEB and Promega), the amount of DNA recovered from muscle was too low for HMW DNA estimation. The Circulomics kit yielded substantially more DNA (> 25 μg for gonad) than other kits and the DNA retrieved had a relatively high proportion of HMW DNA (>90% for gonad). We therefore focused on the Circulomics kit and repeated the Circulomics extractions for an additional four individuals, using the two most promising tissue types: liver and gonad. In line with the first extractions, the additional extractions yielded on average substantially more DNA than the other extraction kits. Among Circulomics DNA extractions, gonad tissues yielded consistently more (mean 33.0 μg vs. mean 15.9 μg) and a higher proportion of HMW DNA (mean 98.4% vs. mean 74.6%) than liver tissues. Across all metrics, for these small lizards, the Circulomics Nanobind animal tissue extraction kit in combination with gonad tissue was the most promising strategy for HMW DNA extraction and LRS. Moreover, this combination also yielded purity measurements that most closely and consistently matched manufacturer’s recommendations for PacBio HiFi sequencing (see Figure 1 for range in grey).

### A high-quality de novo genome assembly for C. pulcher

PacBio sequencing across two 8M SMRTcells yielded 39.5 gigabases (Gb) of adapter filtered HiFi reads with a mean read quality of 27.1 and a read N50 of 16,951 bp, representing ∼27x coverage based on the expected genome-size (1.5 Gb). Using hifiasm, we generated a high-quality primary assembly that included 100 contigs with a contig N50 value of 42.9 Mb. Following duplicate purging, the N50 value remained constant at 42.9 Mb but the number of contigs decreased to 59. Notably, 93.7% of the 1.49 Gb genome was contained in contigs > 10 Mb and 99.9% was captured by contigs > 1 Mb. Thus, PacBio HiFi sequencing alone, using a tissue sample collected and transferred in practical conditions for fieldwork, yielded a highly contiguous assembly.

To place the 59 contigs into empirically informed chromosome models, a rare feat for Scincid lizards, we used a tissue sample that was flash frozen and cryopreserved to generate an Omni-C library for chromosome conformation capture. We scaffolded the duplicated purged PacBio contigs with YaHS and made 1 manual correction, yielding 16 total scaffolds that contained all 59 contigs (Figure S1). After assembling and curating a mitochondrial genome from these same reads, the final genome was 1.49 Gb, and had a scaffold N50 of 199.8 Mb, the length of the 4th longest chromosome model. A BUSCO evaluation indicated that the assembly is highly complete, with 97.9% of vertebrate BUSCOs present and complete (C:97.9%[S:96.9%,D:1.0%],F:0.8%,M:1.3%,n:3354). Based on Omni-C contact density (Figure S1), we found support for 15 chromosome pairs in *C. pulcher*. This is one more than *C. pulcher*’s closest karyotyped relative; *Cryptoblepharus boutonii* (2n=28; [52]). We found evidence of 13 telomeres in the *C. pulcher* genome: two chromosomes were gapless and telomere-to-telomere (T2T), while nine chromosomes had a single telomere. Annotation with RepeatMasker indicated that 47.0% of the genome is classified as repetitive, which is slightly higher than the 44.9% reported for *C. egeriae* [23].

### Genome synteny and sex chromosomes

Using the Omni-C informed chromosome models for *C. pulcher*, we then scaffolded the highly contiguous (contig N50 109.1 Mb; 10 contigs T2T), but still contig-level *C. egeriae* assembly. Reference guided scaffolding made seven joins between *C. egeriae* contigs, resulting in 15 chromosome-models that contain 99.6% of total genome sequence, with the remaining 0.4% of the genome distributed among 50 short contigs (N50 192 kb). Notably, in addition to the ten contigs that were T2T in the unscaffolded *C. egeriae* assembly, reference guided scaffolding with *C. pulcher* resulted in an additional four scaffolds with telomeres on both ends (Figure 2a). Thus, 14 of the 15 *C. egeriae* chromosomes represent telomere-to-telomere contigs or scaffolds, which is remarkable for a non-model and threatened species.

**Figure 2.**
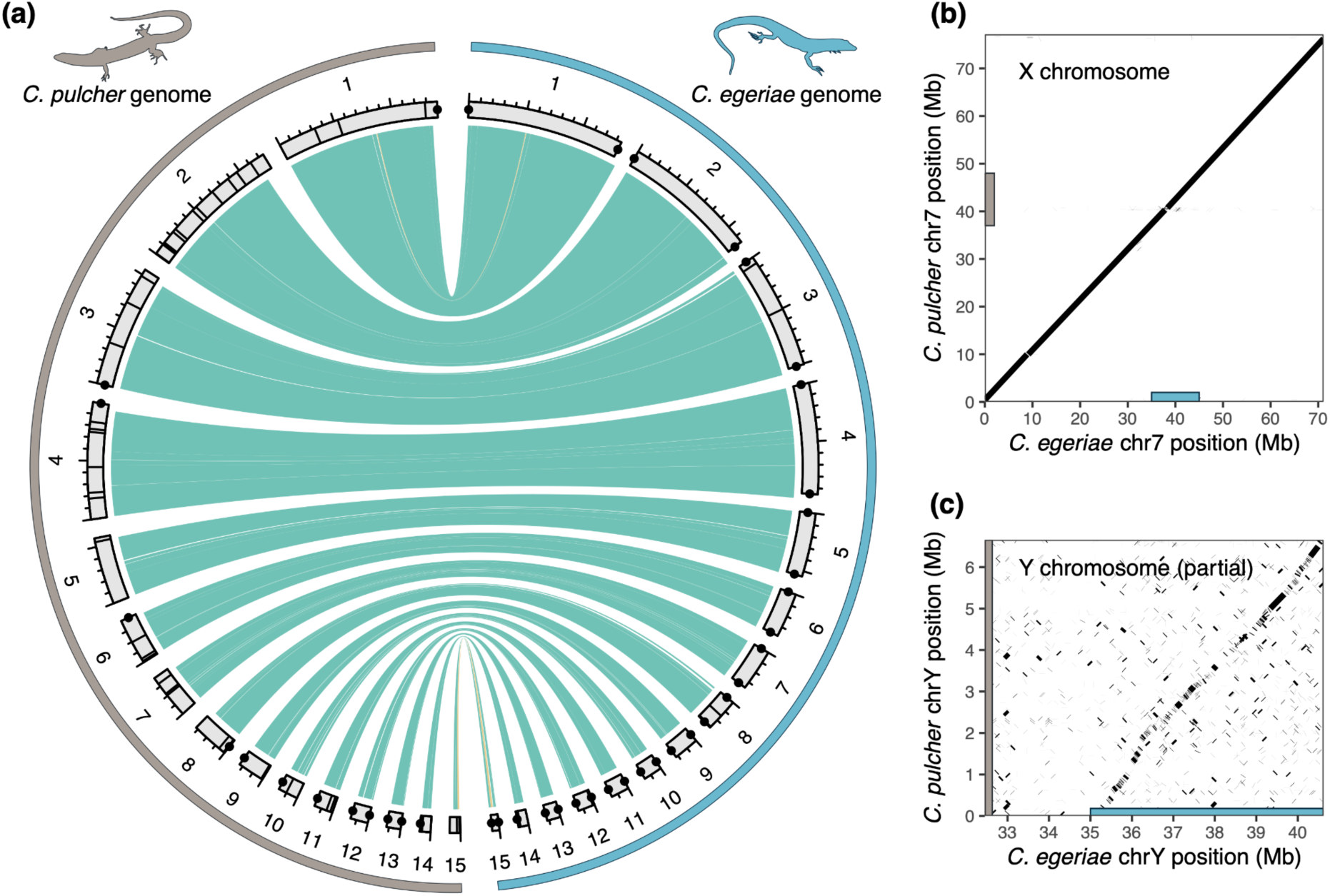
Comparison of genome synteny between *C. pulcher* and *C. egeriae*. (a) *C. pulcher* and *C. egeriae* genomes display low levels of chromosomal rearrangements. Circos plot of minimap2 alignments between *C. pulcher* (left) and *C. egeriae* (right) genome assemblies. Alignment blocks > 1Mb are shown. Co-linear blocks are colored turquoise and inversions are colored beige. Large and small tick marks for each chromosome 100 Mb and 20 Mb, respectively. Telomeres are denoted with black dots and gaps, joined by either Omni-C proximity ligation or by homology, are shown with black lines. Note: *C. egeriae* sequences have been reverse-complemented to aid visualization. (b) Putative X chromosome alignment shows high sequence conservation. Bars on X and Y axes show regions where coverage decreases to half fold, suggesting the sex-linked regions. (c) Putative Y chromosome alignment of the partial sequences shows identity, but also suggests divergence. Bars on X and Y axis same as (b).

Genome alignments between the *C. pulcher* and *C. egeriae* assemblies revealed high levels of synteny. Despite an estimated divergence time of 12 million years between these species [17], when considering structural differences > 1 Mb in the alignments, we only detected two inversions and no inter-chromosomal rearrangements across the entire genome (Figure 2a). A comparison of the putative sex-linked regions revealed good alignment between the putative X chromosomes (Figure 2b), as identified based on coverage (Figure S2). The partial Y chromosome generated by the assembly (see Methods) also showed identity to the *C. egeriae* Y; however, the alignment was substantially more divergent compared to the alignment between X chromosomes (Figure 2c, Figure S3), which may be due to repeat accumulation during Y chromosome degeneration.

### Heterozygosity, historical demography and inbreeding

Previous analyses of the *C. egeriae* genome reported a surprising degree of genetic diversity, large historical population sizes and some signatures of inbreeding [23]. However, the relative magnitude of these estimates remained difficult to appraise without comparative information available for any other skink, let alone congeneric taxa. We explored patterns of heterozygosity across the two *C. pulcher* individuals (one sequenced with PacBio HiFi and a second with Illumina for Omni-C) and repeated the analysis for the new *C. egeriae* chromosome-level assembly. Based on our filtering criteria, 95.2% of the *C. pulcher* reference genome was mappable and callable using HiFi reads and 80.2% was similarly accessible to Omni-C reads. 76.8% of the reference genome was accessible to both data types. In comparison, 98.6% of the *C. egeriae* genome was accessible to variant calling with HiFi reads.

At the genome-wide level, both *C. pulcher* individuals showed similar levels of heterozygosity, with a median of 5.06 and 5.38 heterozygous sites per kb for the individuals sequenced with HiFi (Figure 3a-b) and Omni-C (Figure 3c-d), respectively. This was lower than *C. egeriae* (Figure 3e-f), which had a median heterozygosity of 6.91 heterozygous sites per kb, but also included windows where heterozygosity was at or near 0. Using the same observational approach to identify ROH (see methods), we found only two ROHs in both *C. pulcher* individuals (PacBio HiFi and Omni-C), representing 0.26% and 0.20% of the total genome sequence respectively (Figure 3g). This stands in contrast to our estimate of 9.66% of the *C. egeriae* genome falling within ROH, which is identical to previous estimates using the contig-level assembly [23]. Finally, we used PSMC to infer historical population sizes from these data. In line with high rates of heterozygosity, both species had large historical effective population sizes (Figure 3h). However, consistent with the lower rates of heterozygosity, the inferred population size of *C. pulcher* was continuously smaller over time than the population size of *C. egeriae*. This came as a surprise given that *C. egeriae* has been restricted to Christmas Island (land area of 135 square kilometers) for a long time and diverged from its nearest relatives nearly 6 Mya [53], whereas continental *C. pulcher* has a much larger range [24]. To test if this could be an artifact of different regions of the genome being surveyed, we ran PSMC on the set of variants accessible to both HiFi and Omni-C data. The two inferred population histories were similar to those obtained from the set of unrestricted variants (Figure S4), suggesting that the observed variation in historical population sizes were not caused by any (genomic) sampling bias but instead reflects a biological signal that is shared by both *C. pulcher* individuals.

**Figure 3.**
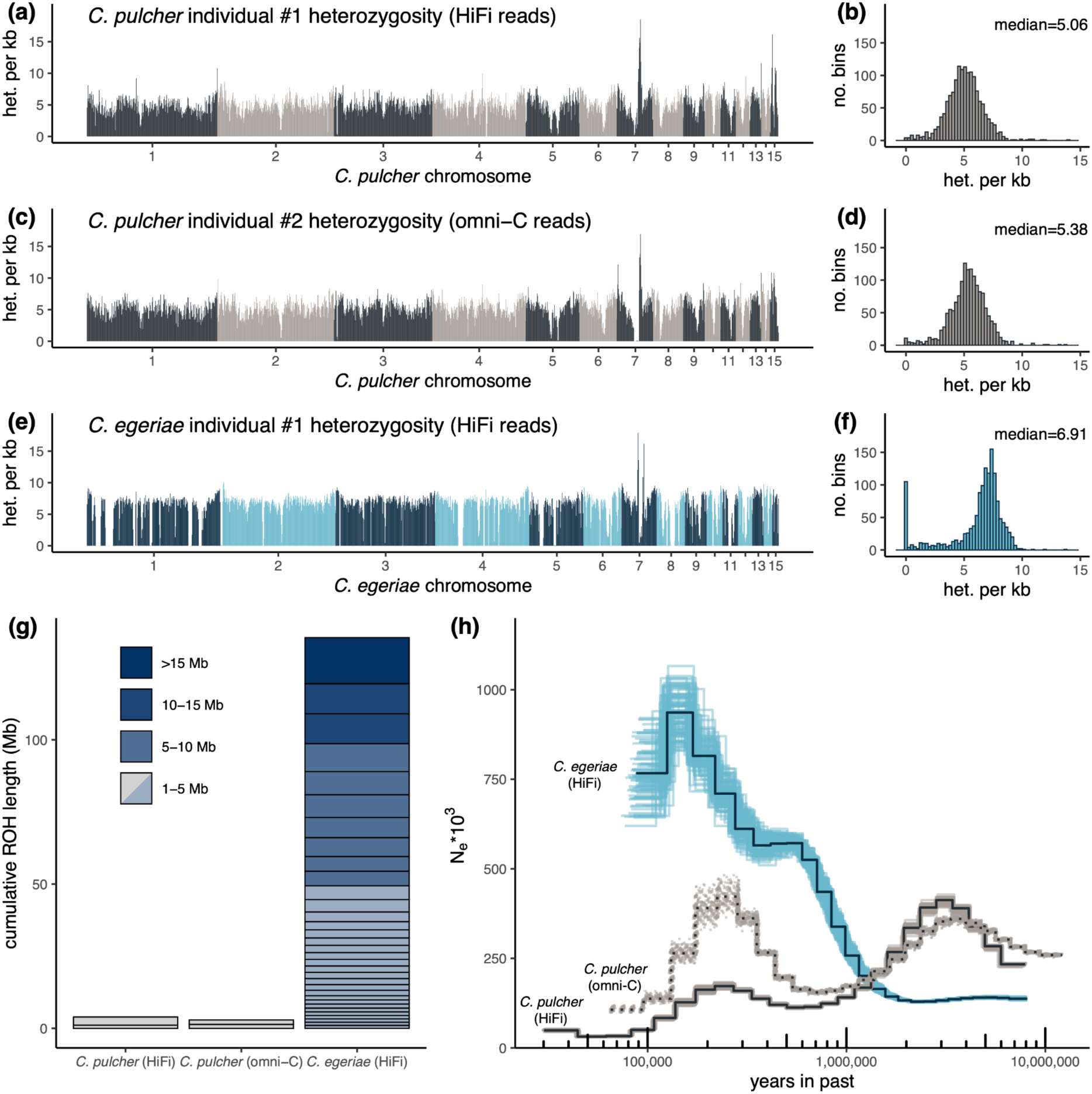
Genome-wide patterns of genetic diversity and demographic history. (a) Heterozygosity across the genome of the first *C. pulcher* individual, sequenced with HiFi reads. Heterozygous SNPs per kilobase (kb) are plotted in 1 Mb bins across the genome, with chromosomes colored in alternating colors. (b) Histogram of windows from *C. pulcher* HiFi data, showing that median heterozygosity falls at 5.06 SNPs per kb. (c) Heterozygosity across the genome of the second *C. pulcher* individual, sequenced with Omni-C reads. (d) Histogram of windows from *C. pulcher* Omni-C data, showing that median heterozygosity falls at 5.38 SNPs per kb. (e) Heterozygosity across the genome of *C. egeriae*, previously sequenced with HiFi reads (Dodge et al. 2025). (f) Histogram of windows from *C. egeriae* HiFi data, showing that median heterozygosity falls at 6.91 SNPs per kb. (g) Cumulative ROH lengths (>1Mb) for each of the three individuals. The two *C. pulcher* individuals sequenced with different technologies each have 2 ROH, covering 0.26% and 0.20% of the genome, while the *C. egeriae* individual has 33 ROH, covering 9.66% of the genome. (h) PSMC plot reflecting the inferred effective population sizes over time for the three individuals sequenced. All individuals are scaled with generation time θ = 3.5 and a median Squamate mutation rate of μ = 6.125 × 10^−9^. Lighter colors denote 100 bootstrap replicates.

## Discussion

Our contemporary and future proficiency to assay the full spectrum of genetic diversity depends on our ability to retrieve high-quality DNA (and other molecules) from samples distributed across the world. However, with biodiversity declining at an unprecedented rate [54], there is an urgent need for cost-effective and practical methods that preserve DNA integrity when sampling, transferring and storing tissues [11]. The objective of the present study was two-fold. First, we wanted to explore the feasibility of using a pragmatic sampling strategy for assembling a high-quality lizard genome. Second, we specifically wanted to generate a genomic resource for a second *Cryptoblepharus* species, *C. pulcher*, to shed more light on the recently reported evolutionary history and corresponding genomic changes of *C. egeriae*, an insular skink that is extinct in its native habitat. Given these broad objectives, further replicating and statistically verifying sampling and extraction strategies was outside the scope of the present study. Rather than providing definite guidelines, our methodological findings should therefore be interpreted as a proof-of-concept sampling and sequencing strategy for Squamate genomics (when gold standard collection guidelines cannot be followed). Most importantly however, we provide an empirical demonstration that it is certainly feasible to generate high-quality genomic resources using practical and cost-efficient sampling strategies. Even without the chromosome-conformation capture data (for which cryopreservation is often needed), the *C. pulcher* genome was already represented in 59 long contigs (N50 = 42.9 Mb). Interestingly, the contig N50 of the *C. egeriae* genome, also assembled with PacBio HiFi reads, was more than double (N50 = 109.1 Mb), despite having a lower HiFi read N50 than *C. pulcher* (12,885 bp vs 16,951 bp respectively). We initially suspected that this might be due to the difference in sequencing depth between both species (∼27x and ∼40x), but down-sampling the *C. egeriae* reads surprisingly did not support this hypothesis (Table S2). While the precise explanation remains elusive, the *C. pulcher* genome already reaches a very high standard and provides an empirical demonstration that it is feasible to generate contiguous assemblies without using costly and impractical sampling strategies.

In line with standardized benchmark tests for other vertebrates [12] and invertebrates [15], we find that lizard tissue samples preserved at ambient temperature in >95% EtOH can still yield HMW DNA more than a week after sampling. Empirical tests are an important validation of standardized experiments, because experiments in controlled settings can sometimes bypass unforeseen factors of importance. For example, ambient temperature may vary more strongly and erratically during shipping than during a controlled experiment in a laboratory. However, besides three extracts, more than 50% of the DNA extracted was > 40 kb in length across all extraction kits and tissue types (Figure 1), suggesting that EtOH with a high purity (>95%) is a promising alternative when cryopreservation is not feasible; at least at short to intermediate transfer/storage times. Notwithstanding these promising results, it is important to emphasize that the focus of the present study was on recovering DNA suitable for *de novo* genome assembly. It remains unclear whether similar sampling strategies are also suitable for preserving other molecules such as RNA or proteins. Furthermore, while it is a promising alternative, suboptimal sampling will certainly reduce the likelihood of obtaining ultra-High Molecular Weight DNA (> 1 Mb fragments) and will therefore limit the potential for generating ultra-long reads [12]. However, we argue that from the perspective of evolutionary and conservation biologists, the development of taxon specific practical sampling strategies, beyond the “gold-standard” of cryopreservation, such as presented here, is of utmost importance [11]. The logistical and financial burden associated with cryopreservation poses a significant challenge for many field biologists and can demotivate efforts to collect high-quality tissue samples, particularly in remote tropical regions that are difficult to access but simultaneously harbor most of the global biodiversity [55]. Practical approaches may be suboptimal, but our study demonstrates that they can still yield contiguous DNA fragments, which could be leveraged to address a wide array of evolutionary and conservation questions. These approaches therefore represent valuable alternatives when cryopreservation is not an option and are particularly relevant considering the increasing demand for high-quality tissue samples (e.g. to construct pangenomes [16]).

Besides investigating to what extent a practical sampling strategy could beget a high-quality assembly, we specifically focused on *C. pulcher* to learn more about genome evolution within skinks and to enable a comparative view on the genetic diversity and demographic history of *C. egeriae*. Scincidae is the largest family among lizards (1,579 species) but there are only six species for which a long-read genome assembly has been released to date (https://www.ncbi.nlm.nih.gov/datasets/genome/?taxon=66056; last accessed August, 1^st^ 2026). With the independent assembly of *C. pulcher*, we are now for the first time able to compare the genomes of two congeneric skink species that are relatively similar in ecology and life-history traits. Even though they have been independently evolving for 12 million years [17], the two genomes exhibit a marked degree of stability in terms of genome synteny; echoing karyotype work done in other skinks [56]. We surveyed the genome for any major rearrangements or structural differences (> 1 Mb) and only identified two inversions (one on Chr1 and another on Chr15; Figure 2). Like the autosomes, the X chromosomes between the two species align closely and show a relatively high degree of conservation (Figure 2b). In contrast, the putative Y chromosomes differ substantially (Figure 2c) and yield patterns that are consistent with a model of rapid Y chromosome degeneration [57]. These findings expand our knowledge of Squamate genome evolution at more recent timescales and complement our existing view on the known dynamism of fissions, fusions and rearrangements of chromosomes throughout reptile evolution [58].

Populations of the Christmas Island blue-tailed skink (*C. egeriae*) started to rapidly disappear by the early 2000’s; two captive breeding populations were established by 2010 and the species was officially declared as Extinct-in-the-Wild by 2017 [20]. The individual selected for *de novo* genome assembly belonged to the captive breeding population at Taronga Zoo (Sydney, Australia) and the high-quality assembly yielded a surprising view on the genetic composition and history of the species [23]. By now sequencing a continental equivalent that is widely distributed, our findings provide a comparative context that sheds further light on the demise and, potentially, future recovery of the species following captive breeding [21]. Both *C. pulcher* individuals (different individuals were used for PacBio HiFi and Omni-C sequencing) have a slightly lower genetic diversity, measured as mean genome-wide heterozygosity. In parallel, the PSMC inference of both individuals suggests that on average the effective population size of *C. pulcher* was lower than the population size of *C. egeriae* throughout the past 1 million years (Figure 3). These findings are surprising because the Christmas Island population has been genetically isolated from other extant *Cryptoblepharus* species for millions of years, on an island that is roughly 135 km^2^ in size. In contrast, *C. pulcher* is widely distributed in coastal woodland habitat across most of eastern Australia. Notwithstanding this widespread distribution, it could be that the *C. pulcher* population sampled near Brisbane is relatively isolated or underwent a recent bottleneck, but there are no obvious indications that support either hypothesis. Alternatively, it could be that the *C. egeriae* population on Christmas Island experienced less competition and/or native predation than their continental equivalent, leading to a relatively high population density in a much more confined area. The latter hypothesis aligns with surveys conducted before the introduction of invasive species and personal observations up until at least 1990, which reported them as ‘hyper abundant’ in the island’s settlement area [59].

When taking genome-wide heterozygosity as a proxy for the genetic diversity of both species, our comparative estimates suggest that the initial captive breeding population of 66 *C. egeriae* individuals included a substantial fraction of the genetic diversity segregating in the original population; at least to the same level as in a continental and ecological analogue. This finding supports a model where the demise of *C. egeriae* was primarily caused by invasive predators that rapidly drove the entire Christmas Island population to extinction [60] and lends less weight to a possible ‘extinction vortex’ scenario. Furthermore, since the *C. pulcher* population is seemingly healthy at the same level of genetic diversity, the current *C. egeriae* breeding population should harbor sufficient diversity to establish new populations in the wild. Trial introductions of captive bred individuals in both its native range [61] and on an isolated extralimital atoll [22] have shown promising signs that further support this notion. However, it is important to recognize that genome-wide heterozygosity is merely one indicator of genetic diversity and captive breeding can artificially inflate heterozygosity. This can sometimes even lead to outbreeding depression, but this is not likely for *C. egeriae* since all individuals sourced for the captive breeding program were caught in a relatively small geographic area and belonged to the same island population [21]. Yet, with this data, we cannot exclude the possibility that adaptive alleles were lost and some signs of inbreeding are detectable in the *C. egeriae* genome that are absent in *C. pulcher* (i.e. F_ROH_, 0.0966 vs. 0.0025 respectively). Future studies should investigate to what extent these ROHs may have led to Loss-of-Function substitutions and how fast these ROHs can be broken-up with further assisted breeding or in the wild [62].

In summary, our findings demonstrate that a pragmatic sampling strategy can still yield high-quality genomic resources and provide the foundation for comparative studies that assay evolutionary and genomic change. At the same time, our findings also highlight that the quality, quantity and yield, can vary between tissue types and extraction kits. It is important to note that this may be taxon specific and should encourage broader exploration across organismal groups. Moreover, while we demonstrate that ethanol-preserved tissues can still be used to assemble a high-quality vertebrate genome, more research is needed to explore whether chromatin conformation capture can also be done without flash-frozen tissues. A recent study [15] in an invertebrate (mosquito) presented a protocol which allows the use of nuclei stored for ∼1 week at ambient temperatures, and a similar approach could also be tested in vertebrates. With biodiversity declining at a rapid rate, advances in sampling protocols across the Tree-of-Life will become increasingly relevant for molecular ecology, conservation science and evolutionary biology.

## Acknowledgements

The authors acknowledge support from the National Genomics Infrastructure in Stockholm funded by Science for Life Laboratory, the Knut and Alice Wallenberg Foundation and the Swedish Research Council, and SNIC/Uppsala Multidisciplinary Center for Advanced Computational Science for assistance with massively parallel sequencing and access to the UPPMAX computational infrastructure. We would like to particularly acknowledge Martin Irestedt, Olga Vinnere Pettersson and Elísabet Einarsdóttir for their advice and help in generating the data. Finally, we are very grateful to Saphira Bloom-Quinn for her assistance in transferring samples from Australia to Europe. TOD was supported by NSF GRFP award DGE-2146755. This work was funded by the Innovation Fund of the Museum für Naturkunde Berlin (grant awarded to MPKB).

## Data Accessibility and Benefit-Sharing

### Data Accessibility

Raw sequencing reads and assemblies are all publicly available from NCBI SRA under BioProject numbers PRJNA1287626 (*C. pulcher*) and PRJNA924831 (*C. egeriae*). The code underlying analyses is available at https://github.com/tododge/cpulcher_assembly.

### Benefit Sharing

A research collaboration was established between the Museum für Naturkunde Berlin (Germany) and the Queensland Museum in Brisbane (Australia); all collaborators have been included as coauthors. The study explicitly contributes to the overall objective of Benefit Sharing in multiple ways. First, it presents a cost-effective alternative to optimally preserve DNA integrity and thereby lowers the barrier for safeguarding biodiversity genomic resources. Second, it provides a genetic framework to better understand the causes and consequences underlying the population demise of an Australian endemic vertebrate. These insights have been shared with Parks Australia prior to publication. Finally, the high-quality reference genome of *Cryptoblepharus pulcher* has been publicly released and represents an important new resource that facilitates further investigation into the species diversity of Australian skinks.

## Ethics Statement

Animals were collected with permission of the Queensland Department of Environment Heritage and Protection permits WIMU17395016 and under Queensland Museum Animal Ethics committee approvals AEC18-02 and AEC21-02.

## Author contributions

Conceptualization: MPKB, PO

Methodology: MPKB, ME, TOD, PO

Investigation: MPKB, ME, TOD, PO

Visualization: ME, TOD

Funding acquisition: MPKB

Project administration: MPKB

Supervision: MPKB

Writing – original draft: MPKB

Writing – review & editing: MPKB, ME, TOD, PO

