## Supplementary Information for "A practical sampling strategy for biodiversity genomics of reptiles: The *Cryptoblepharus pulcher* assembly offers insights into the demise of a threatened relative"

### Supplementary Figures & Tables:

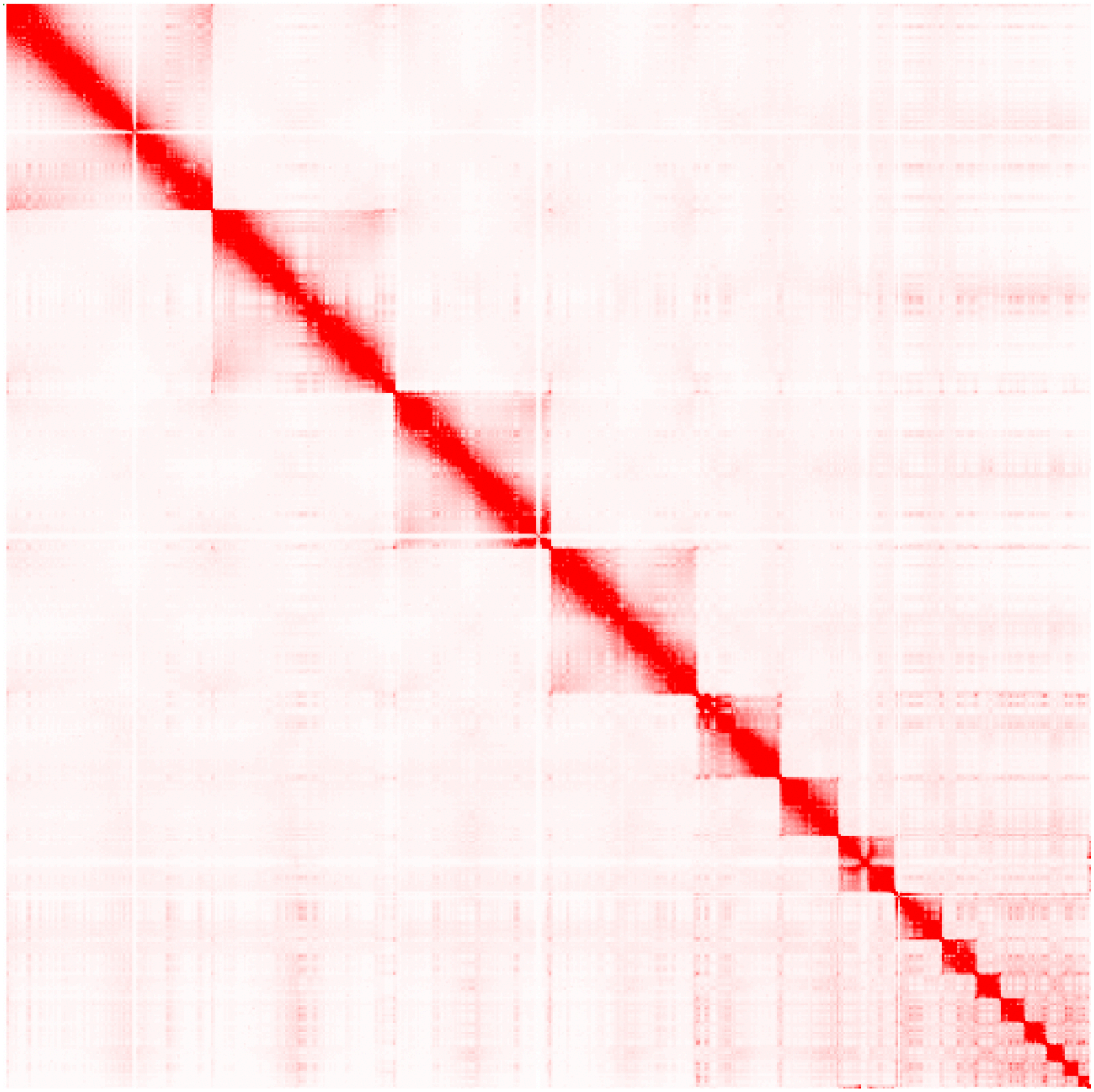

**Supplementary figure 1. Omni-C contact density map.** Chromatin conformation capture data of a *C. pulcher* tissue provides strong support for 15 chromosome models.

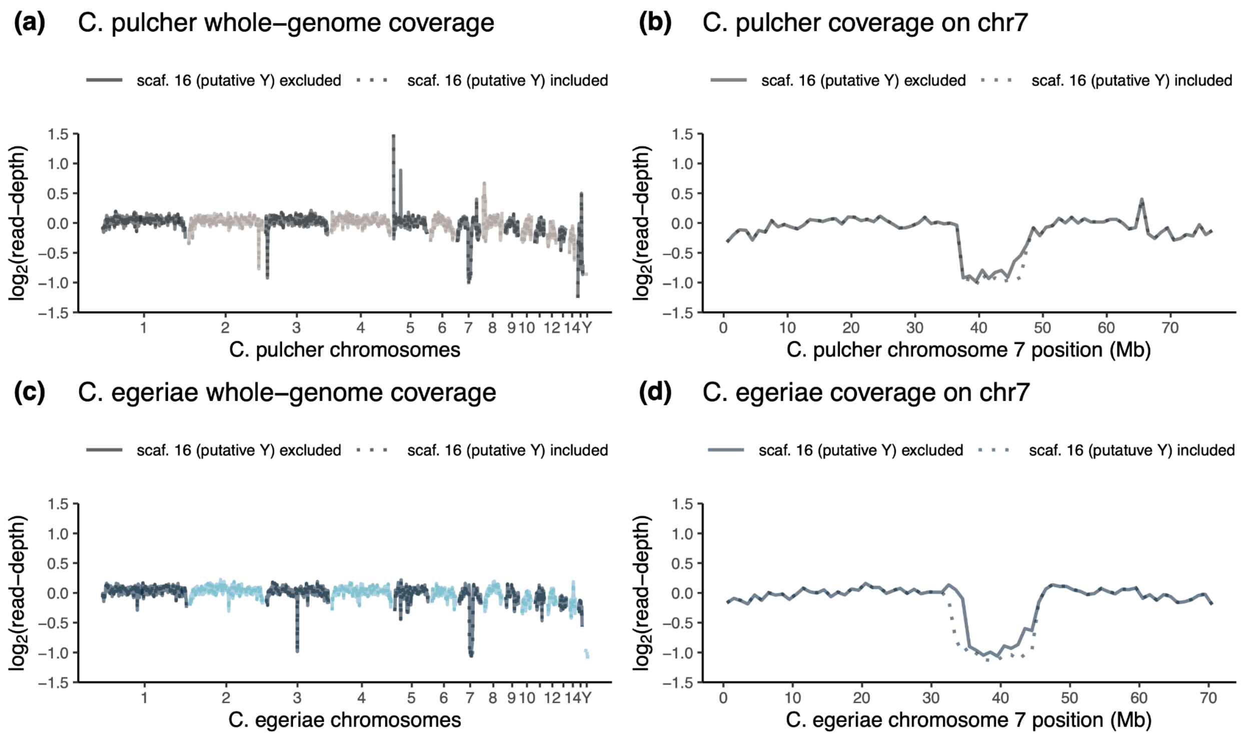


**Supplementary figure 2. Genome-wide coverage comparisons of *C. pulcher* and *C. egeriae*.** Scaled genome wide coverage plots of both genomes, with a focus on the change in coverage when including or excluding the putative Y chromosome. Excluding the putative Y chromosome (scaffold 16) does not change coverage much across most of the genome for either species (a, c), except for two regions, one being the putative pseudoautosomal region of scaffold 7 (b, d). This provides further support for the notion that scaffold 7 likely represents chromosome X and scaffold 16 a partially assembled Y chromosome.

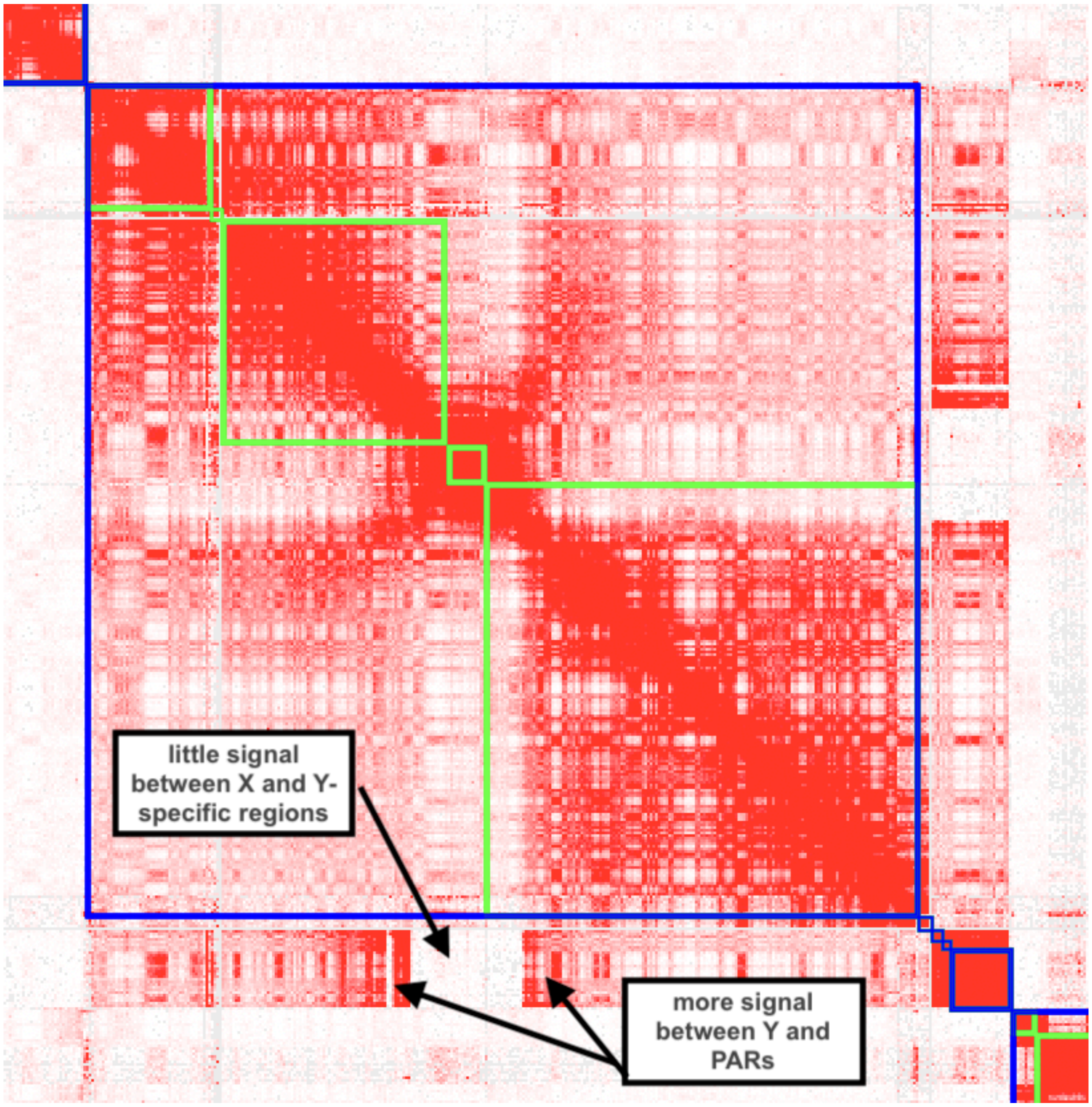


**Supplementary figure 3. Hi-C contact density map between scaffold 7 and 16.** Chromatin conformation capture data show strong affinity between scaffold 7 and 16, with the sequences flanking the ½ coverage region in the middle of scaffold 7 (i.e. the putative pseudoautosomal regions of scaffold 7) but not with the X and Y specific regions.

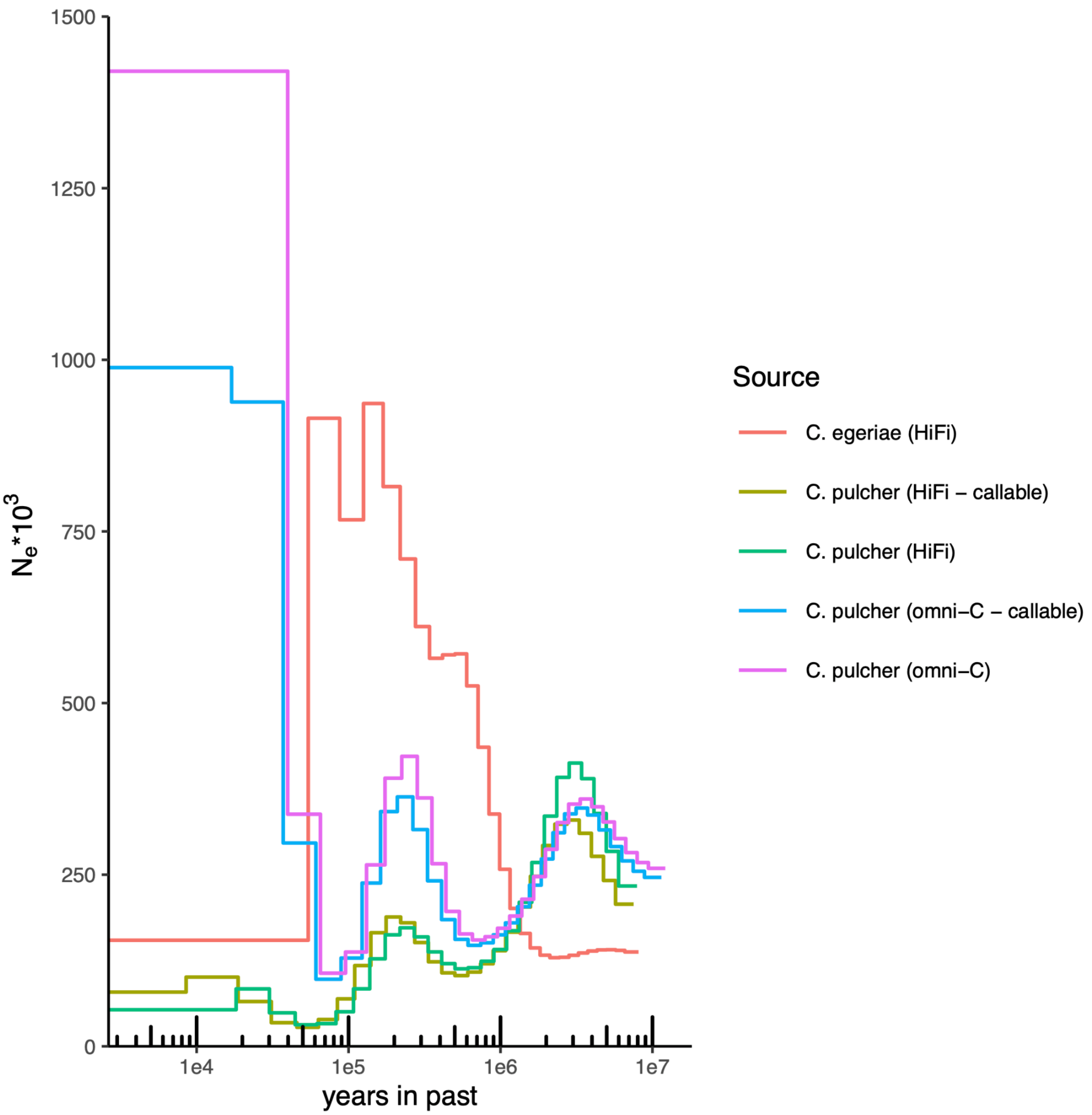


**Supplementary figure 4. PSMC plots for *C. pulcher* and *C.egeriae.*** Individual PSMC analyses were inferred based on distinct datasets for the *C. pulcher* data. The Pacbio HiFi data and Illumina short-read data (omni-C) were split in two categories each, the entire dataset and the region of the genome that were accessible to both the Pacbio and Illumina data.

**Table 1.** Overview of extraction kits and their pricing (at time of study)

| Kit | Manufacturer | Extraction Method | Catalog Number | Price ($) | Number of Reactions | Price per Reaction ($) |
|---|---|---|---|---|---|---|
| Nanobind Tissue Big DNA Kit | Circulomics | Magnetic Disks | NB-900-701-01 | 300 | 24 | 12.5 |
| Wizard HMW DNA Extraction Kit | Promega | Precipitation | A2920 | 343 | 50 | 6.9 |
| MagAttract HMW DNA Kit | Qiagen | Magnetic Beads | 67563 | 427 | 48 | 8.9 |
| Monarch HMW DNA Extraction Kit | New England Biolabs | Glass Beads | T3060S | 74 | 5 | 14.8 |

**Table 2.** Unpurged assembly contiguity and downsampling strategy

| species | assembly (strategy) | raw data (coverage) | read N50 | contig N50 | # contigs | reference |
|---|---|---|---|---|---|---|
| *C. pulcher* | full dataset | 39.5 Gb (27x) | 16951 | 42.9 Mb | 100 | this work |
| *C. egeriae* | downsample (random) | 39.3 Gb (27x) | 12887 | 100.8 Mb | 114 | this work |
| *C. egeriae* | downsample (keep longest reads) | 39.5 Gb (27x) | 14364 | 108.7 Mb | 105 | this work |
| *C. egeriae* | full dataset | 58.6 Gb (40x) | 12885 | 109.1 Mb | 76 | Dodge et al. (2025) |
